# Exercise induces anti-inflammatory reprogramming in macrophages via Hsp60

**DOI:** 10.1101/2025.03.03.640779

**Authors:** Antonios Chatzigeorgiou, Athanasios Moustogiannis, Panagiotis F. Christopoulos, Nikolaos I. Vlachogiannis, Filippos Michopoulos, Kleio-Maria Verrou, Grigorios Papadopoulos, Rallia-Iliana Velliou, Eirini Giannousi, Ioannis Mitroulis, Jindrich Chmelar, Theodossis A. Theodossiou, Martina Samiotaki, Anastasios Philippou, Petros P. Sfikakis, Michael Koutsilieris

**Affiliations:** Department of Physiology, Medical School, National and Kapodistrian University of Athens, 75 Mikras Asias, 11527 Athens, Greece; Department of Pathology, Section of Research, Rikshospitalet, Oslo University Hospital and University of Oslo, Oslo, Norway; First Department of Propaedeutic Internal Medicine, National and Kapodistrian University of Athens Medical School, Athens, Greece; Bioscience, Early Oncology R&D, AstraZeneca, Cambridge, UK; Center of New Biotechnologies & Precision Medicine, National and Kapodistrian University of Athens Medical School, Athens, Greece; First Department of Internal Medicine, University Hospital of Alexandroupolis Democritus University of Thrace Alexandroupolis Greece; Faculty of Science, University of South Bohemia in České Budějovice, 37005 České Budějovice, Czech Republic; Department of Radiation Biology, Institute for Cancer Research, Oslo University Hospital, Oslo, Norway; Institute for Bioinnovation, Biomedical Sciences Research Center ‘Alexander Fleming’, Vari, Greece

**Author notes:** Correspondence to: (Lead contact): Assoc. Prof. Antonios Chatzigeorgiou, MD, PhD, Department of Physiology, Medical School, National and Kapodistrian University of Athens, 75 Mikras Asias Str., 11527, Athens, Greece., and Professor Michael Koutsilieris, Department of Physiology, Medical School, National and Kapodistrian University of Athens, 75 Mikras Asias Str., 11527, Athens, Greece. These authors contributed equally to this work (Antonios Chatzigeorgiou, Athanasios Moustogiannis).

**Keywords:** exercise, Hsp60, skeletal muscle cells, macrophages, inflammation, oxidative phosphorylation, -omics

## Abstract

Physical activity exerts systemic anti-inflammatory effects and reduces the risk for multiple non-communicable diseases, with 7.2% of all-cause deaths globally being attributed to physical inactivity. However, the cellular and molecular components of the exercise-induced anti-inflammatory effects remain only partly understood. Herein we show that moderate-intensity exercise promotes anti-inflammatory reprograming of macrophages orchestrated by the skeletal muscle cells secretome. Primary bone marrow-derived macrophages (BMDMs) exposed to the secretome of mechanically-loaded myotubes (exercise-conditioned medium, exCM) acquire an anti-inflammatory transcriptional profile and increased reliance on oxidative phosphorylation, as shown by Seahorse real-time cell metabolic analysis, compatible with an M2-like phenotypic switch. Using an unbiased proteomic analysis of the exCM we identify the chaperonin Hsp60 as a key mediator of the anti-inflammatory effects of exercise. Hsp60 expression increases in mechanically loaded myotubes in vitro, in the quadriceps muscle and serum of mice following an 8-week program of moderate-intensity aerobic exercise, as well as in human muscle after resistance training. Importantly, treatment of BMDMs with Hsp60 in vitro recapitulates the exCM-induced transcriptional reprograming, promoting an M2-like phenotype. Taken together, our data highlight Hsp60 as a novel component of the skeletal muscle cell-macrophage crosstalk, providing mechanistic insights into the anti-inflammatory effects of exercise.

## Introduction

Macrophages are innate immune cells with a central role in tissue homeostasis and remodeling, originating either from embryonic stem cells that are seeded in the tissues before birth or from infiltrating bone-marrow derived monocytes (BMDMs) (1). Based on the tissue microenvironment, macrophages adapt their transcriptional profile and function showing remarkable plasticity, and can either promote local proinflammatory responses or acquire a pro-resolving phenotype (2). Traditionally, the functional heterogeneity of macrophages is represented by a binary model, whereby M1 macrophages show proinflammatory and M2 macrophages anti-inflammatory properties, respectively (3). In addition to the traditional immunoregulatory stimuli (e.g., cytokines), it has recently been shown that changes in cellular metabolism can affect macrophage polarization and function (4–6). Increasing evidence supports the central role of persistent proinflammatory tissue macrophages in the pathogenesis of immune-mediated diseases (7), while identification of the mechanisms driving their phenotypic shift towards the pro-resolving type (M2-like) has gained interest as a therapeutic approach in inflammatory disorders (8).

Physical activity/exercise has been long known as one of the most essential non-pharmacological interventions in the prevention and management of numerous non-communicable diseases (9). It is estimated that 7.2% of all-cause deaths globally may be attributed to physical inactivity (9). However, the cellular and molecular effects of exercise regulating tissue homeostasis remain largely unknown. Exercise can affect the tissue microenvironment by regulating local immune responses and more specifically the status of macrophages (10). While macrophages in obese individuals are skewed towards a proinflammatory M1 phenotype (11, 12), exercise training has been shown to reduce systemic inflammation, probably via favoring an M1 to M2 switch (10, 13), as well as via inhibiting BMDM infiltration into adipose tissue (10, 14). However, the molecular mechanisms by which exercise mitigates the macrophage-mediated inflammation remain uncertain.

Herein, we investigated whether moderate exercise can affect the secretome of muscle cells and in turn induce an anti-inflammatory profile in macrophages in a paracrine/endocrine manner.

## Results

### The secretome of mechanically-loaded myotubes induces anti-inflammatory transcriptional reprograming of macrophages

First, we examined whether moderate intensity exercise affects the cross-communication between skeletal muscle cells and macrophages. Given the technical challenges that prevent the study of paracrine/endocrine responses of muscle cells to mechanical loading *in vivo*, we utilized a cell stretching device that simulates the mechanical stimuli *in vitro* (15, 16). Mechanical loading of differentiated myotubes *in vitro* may mimic exercise-induced loading patterns of skeletal muscle *in vivo*. Given the complexity of the *in vivo* adaptations in response to exercise-induced external loading, *in vitro* models of mechanical loading applied on muscle cells are crucial for understanding the cellular and molecular mechanisms that mediate loading-induced adaptations. By scrutinizing the components of mechanical loading applied on myotubes *in vitro*, we have previously characterized an overall beneficial, muscle-targeted, moderate exercise mimetics protocol (17, 18). Specifically, herein, differentiated C2C12 myotubes were subjected to a 2% elongation protocol for 12h using a cell stretcher, and the culture supernatant was used to treat primary BMDMs (**Figure 1A**). Primary macrophages incubated with the mechanical loading-induced secretome (diluted 1:1 in culture medium, thereafter called exercise-conditioned medium, exCM) for 24 hours displayed enhanced expression of anti-inflammatory genes (*Il10*, *Il13*, *Tgfb1*, *Pparg*, *Arg1* and *Mrc1*(CD206); **Figure 1B**), while proinflammatory gene expression was decreased (*Il1b*, *Nos2, Tnf*, and *Ifng*; **Figure 1C**). These results suggest that the secretome of the loaded myotubes results in a transcriptional reprogramming of BMDMs that resembles the M2-phenotype (19).

**Figure 1.**
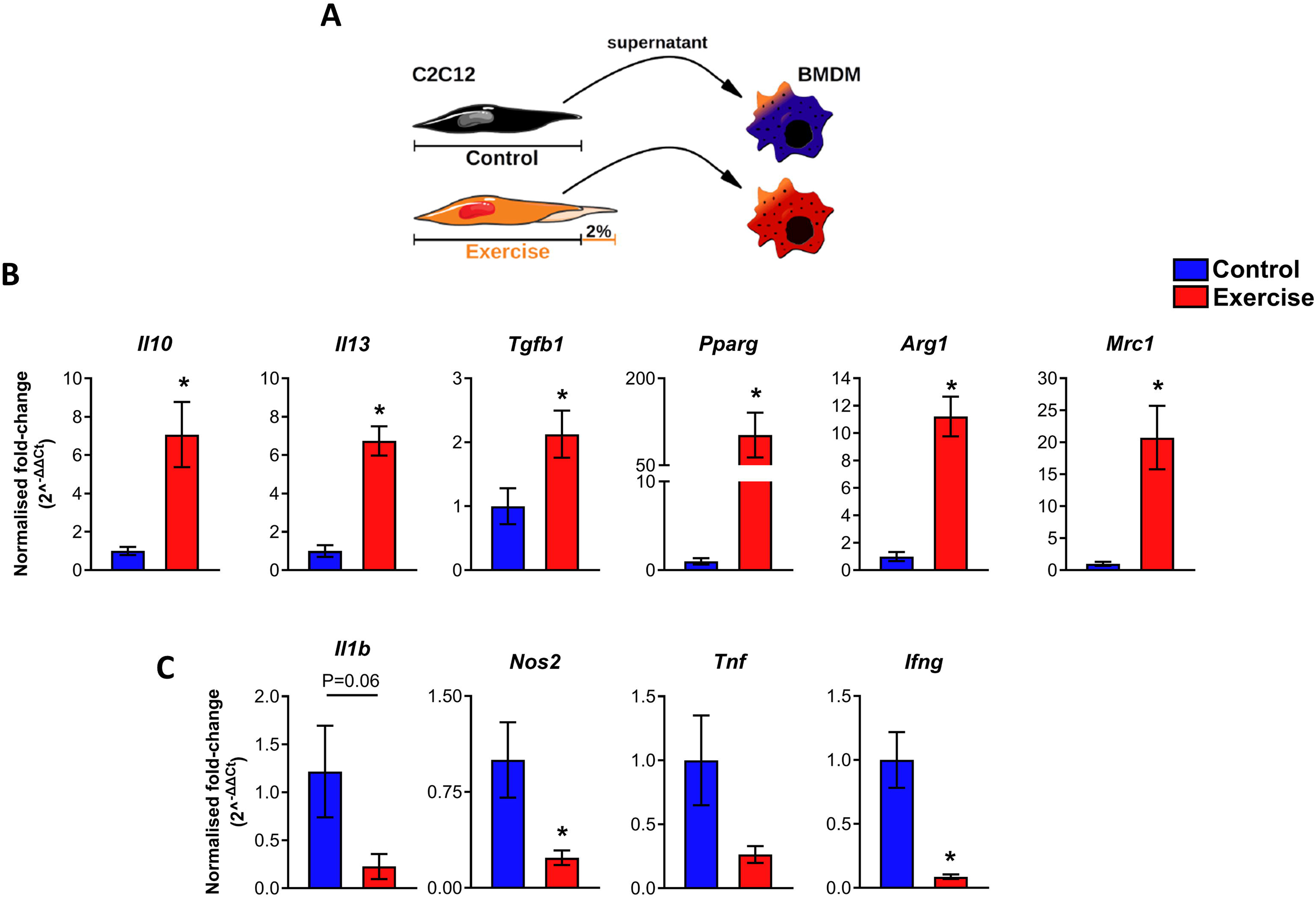
Secretome from stretched myotubes induces an anti-inflammatory phenotype in macrophages. **A)** Fully differentiated C2C12 cells were subjected to 2% elongation for 12h using a cell stretcher. BMDMs were treated with conditioned media (1:1 diluted in culture medium) from control (non-exercised) or stretched (exercise conditioned medium, exCM) myotubes for 24h. **B,C)** qRT-PCR analysis of BMDMs treated with control medium or exCM for quantification of (B) anti-inflammatory or (C) proinflammatory gene expression (n = separate cell isolations from 5 mice). Relative gene expression for each sample is expressed as fold-change *vs* the average value of the control group. Bars represent mean ± standard error of the mean (SEM). Two-tailed Mann Whitney U-test was used to calculate significance. * P˂0.05.

### BMDMs exposed to the secretome of exercised myotubes display increased reliance on oxidative phosphorylation

Recent experimental evidence suggests that metabolic reprogramming is a key factor underlying M1/M2 macrophage polarization (6, 20). In principle, the proinflammatory M1 macrophages rely on glycolysis to cover their energy needs, while the anti-inflammatory M2 macrophages rely mainly on oxidative phosphorylation (21). Having observed that the secretome of moderately loaded myotubes promotes a shift towards an M2-like phenotype in BMDMs, we next examined whether this phenotypic shift is accompanied by metabolic rewiring, performing metabolomic analyses. Indeed, exCM-treated macrophages had increased intracellular levels of tricarboxylic acid (TCA) cycle intermediates, including citric acid (citrate) and a-ketoglutaric acid to support energy flux to oxidative phosphorylation; **Figure 2A** and **Supplementary Figure 1**). The increased demand for nicotinamide adenine dinucleotide (NAD) coenzyme, due to increased use of TCA, was reflected in increased tryptophan metabolism, as tryptophan degradation through the kynurenine pathway is a major source of NAD, ultimately leading to decreased kynurenine levels due to their consumption (**Figure 2A, 2B** and **Supplementary Figure 1**). Increased NADH levels in the exercise group, generated from TCA, provide the necessary electrons to the electron transport chain, leading to increased oxidative phosphorylation (OXPHOS) and ATP production (**Figure 2A** and **Supplementary Figure 1**). Along this line, KEGG pathway analysis of the exCM-treated macrophage metabolome revealed upregulation of processes such as amino sugar, tryptophan and glutamate metabolism, implying a metabolic pattern akin to alternative activation of macrophages (**Figure 2B**). This shift towards increased TCA cycle use and oxidative phosphorylation is compatible with the metabolic signature of M2-like macrophages (21), in line with the transcriptomic rewiring of exCM-treated macrophages presented in Figure 1.

**Figure 2.**
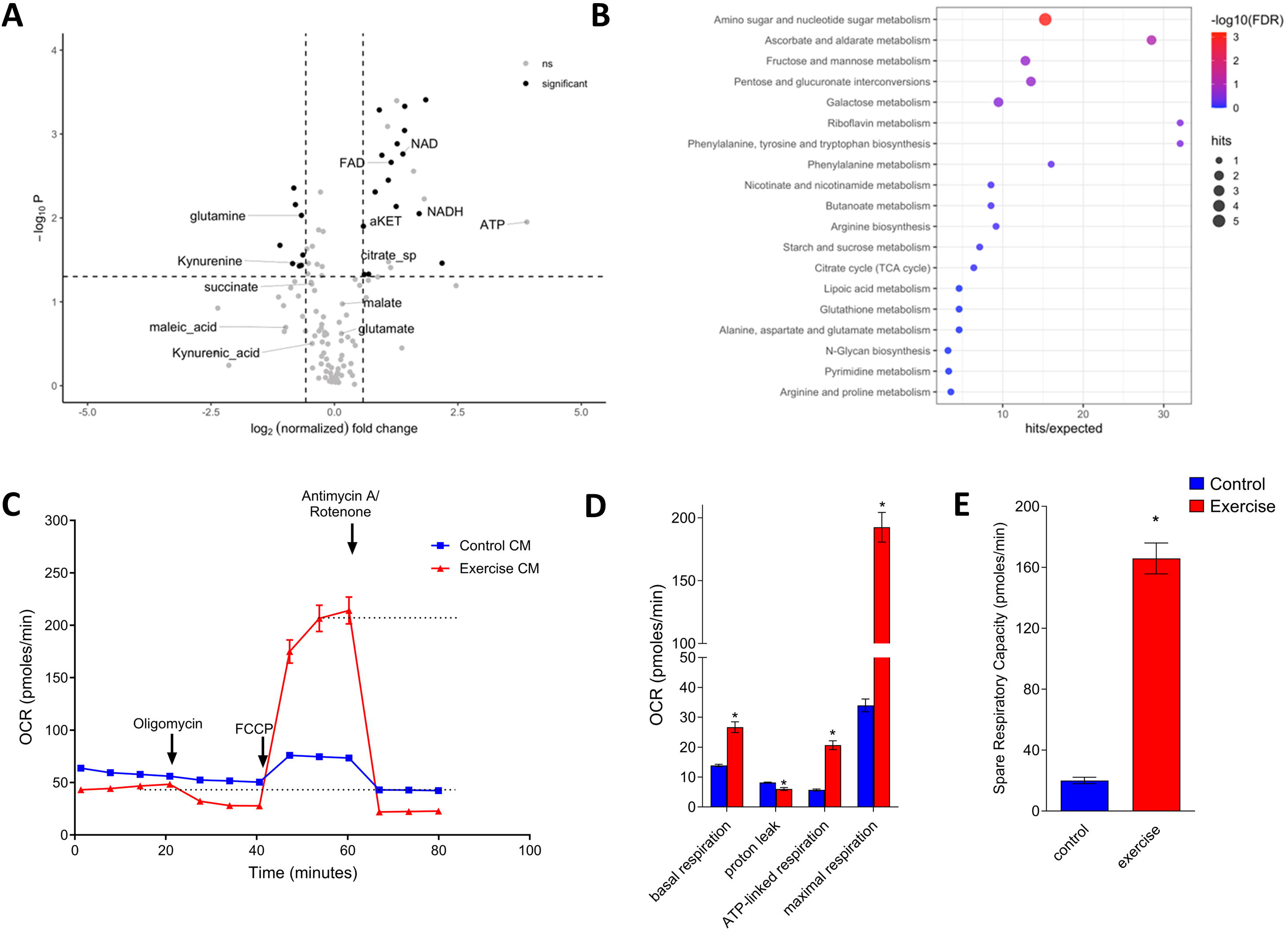
Macrophages treated with secretome from exercised myotubes have enhanced oxidative phosphorylation and increased sugar metabolism. **A)** Volcano plot of differentially expressed metabolites in BMDMs treated for 24 hours with the secretome of exercised *vs* non-exercised myotubes (νon-exercised- vs exercised-treated BMDM from separate cell isolations from n=4 mice). Differentially expressed metabolites are shown with bold (p-value<0.05, absolute log_2_FC>0.58 and covariance <30%). Immune related metabolites involved in the TCA cycle are denoted. **B)** KEGGs pathways’ enrichment plot of overexpressed metabolites (p-value <0.05 and log_2_FC > 0.58) in BMDMs treated with exCM. The enriched pathways are sorted based on FDR, top 3 metabolite pathways have FDR<0.20. **C-E)** Seahorse analysis of BMDMs for respiration (OCR; oxygen consumption rate) bioenergetics. One representative experiment (out of 2 independent) with 14 technical replicates per condition is shown in the line graphs. Bars represent mean ± SEM. Two-tailed Mann Whitney U-test was used to calculate significance. * P˂0.05.

To validate the metabolomic findings, we explored the bioenergetics of exCM-stimulated macrophages using the Agilent Seahorse XF Cell Mito Stress Test, which allows live cell monitoring of the oxygen consumption rate. We separately evaluated the mitochondrial and glycolytic function of cells using the respective Seahorse assays. Extracellular flux analysis confirmed the upregulation of OXPHOS in the exCM-treated macrophages (**Figure 2C**), as depicted in increased basal respiration, ATP-linked respiration, and maximal respiration (**Figure 2D**), as well as by increased spare respiratory capacity following FCCP injection (**Figure 2E**). On the contrary, slight differences were observed in glycolysis between control and exCM-treated macrophages (**Supplementary Figure 2**), despite reaching significance. Taken together, these results show that macrophages treated with exCM acquire a metabolic profile dependent mainly on respiration for their energetic demands, similar to that of M2 macrophages.

Next, to examine whether the observed transcriptomic-phenotypic switch induced in exCM-treated macrophages was driven by increased OXPHOS, we used a selective OXPHOS inhibitor that selectively impairs ETC complex I (IACS-010759) currently used in clinical trials (22). Pre-treatment of BMDMs with IACS-010759 did not affect the anti-inflammatory gene expression induced by exCM (**Supplementary Figure 3A**), while it resulted in enhanced expression of *Tnfa* (p˂0.05) (**Supplementary Figure 3B**), suggesting that upregulation of OXPHOS in exCM-treated macrophages is a core molecular event taking place during their exercise-related reprogramming.

### Unbiased proteomic analysis of exCM identifies Hsp60 as potential mediator of exercise-induced anti-inflammatory effects

Having observed that the secretome from loaded myotubes induces an anti-inflammatory programming in macrophages in a paracrine manner, we sought to identify which secreted factor(s) may mediate this effect. Proteomic analysis of the exCM showed that in total 266 proteins were differentially expressed / secreted upon exercise (**Figure 3A**). Of interest, 52 proteins were related to immune system function (**Figure 3B**), while Hsp60 (Hspd1) stood out as the single protein directly associated with regulation of macrophage activation (**Figure 3C**).

**Figure 3.**
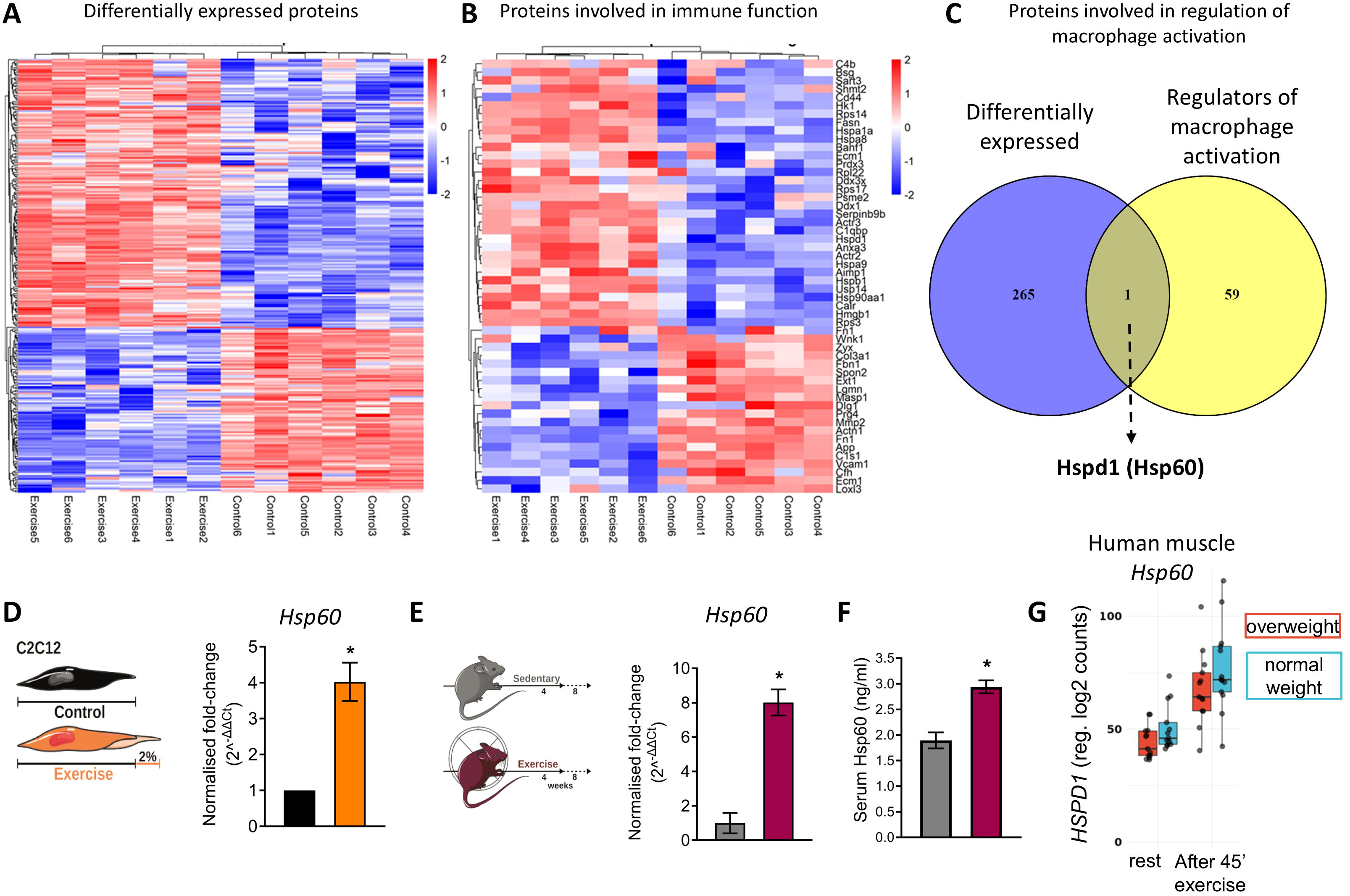
Mechanical loading alters the immunomodulatory secretome of C2C12 myotubes. Fully differentiated C2C12 cells were subjected to 2% elongation for 12h using a cell stretcher. The secretome of exercised and non-exercised (control) myotubes was subjected to proteomic analysis. **A)** Heatmap of Differentially Expressed Proteins (P-adj.<0.05) in exercised and control C2C12 cells. (n= 6 replicates per condition in cell culture). **B)** Heatmap of Differentially Expressed Proteins related to immune response (n=52, with P-adj.<0.05 and log2FC >0.1) **C)** Venn diagram depicting the overlap of the proteins that were differentially expressed in the exercised myotubes with proteins related to “regulation of macrophage activation” (GO:0043030). A single protein (Hspd1/Hsp60) was identified. **D)** *Hsp60* (*Hspd1*) expression (mRNA) in exercised *vs* control myoblasts. Bars represent mean ± standard error of the mean (SEM) of five (n=5) independent experiments performed in duplicate. **E)** Mice were subjected to a moderate intensity exercise program for 8 weeks using running wheels (exercise group, n=5 mice) or not (control, n=5 mice). Bars show mRNA expression of *Hsp60* (*Hspd1*) in isolated quadriceps muscle. **F)** Quantification of serum Hsp60 levels by ELISA in mice subjected to a moderate-intensity exercise program for 4 weeks (exercise group, n=5 mice) or not (control group, n=5 mice). **G)** *HSPD1* (HSP60) expression levels in the *Vastus lateralis* muscle of both normal-weight and overweight individuals following 45 minutes of exercise (re-analysis of MyoGlu dataset; available at https://exchmdpmg.medsch.ucla.edu/app/). In all bar graphs, lines represent the mean and SEM of each depicted group. Two-tailed Mann Whitney U-test was used to calculate significance. * P˂0.05.

### Hsp60 expression is induced by exercise in the muscles of mice and humans

To confirm the proteomic findings, we first tested the expression of *Hsp60* in stretched C2C12 myotubes and verified its upregulation *in vitro* (**Figure 3D**). Next, we examined whether Hsp60 expression was increased in muscle tissue *in vivo* following an 8-week protocol of moderate-intensity exercise in mice (**Figure 3E**). Indeed, analysis of mouse quadriceps muscles revealed the upregulation of *Hsp60* mRNA expression upon mechanical loading *in vivo* (**Figure 3E**). Moreover, increased circulating Hsp60 was detected in the serum of mice following the 8-week exercise protocol **(Figure 3F**), confirming its secretion and potential to exert paracrine/endocrine actions. Of interest, increased expression of *HSPD1* (HSP60) was also observed in human *Vastus lateralis* muscle samples derived from both normal-weight and overweight individuals following 45 minutes of exercise (**Figure 3G**), supporting the relevance of our findings in the human body.

### Hsp60 induces anti-inflammatory transcriptional reprogramming of BMDMs

Finally, we examined whether Hsp60 could recapitulate, at least in part, the effect of exCM treatment on myotubes, thus comprising a key mediator of the paracrine anti-inflammatory effects of exercised myotubes on BMDMs. Extracellular Hsp60 may have a broad spectrum of functions depending on the tissue or cell context (23, 24). To examine the effect of Hsp60 on macrophage phenotypic switch, we treated BMDMs with Hsp60 *in vitro*. Hsp60 treatment resulted in enhanced expression of anti-inflammatory genes (**Figure 4A**) and decreased expression of the pro-inflammatory genes *Il1b, Nos2, Tnfa*, and *Ifng* (**Figure 4B**), recapitulating the anti-inflammatory shift induced by exCM treatment of macrophages (Figure 1), and suggesting that Hsp60 is indeed a main driver of the macrophage phenotypic switch induced by exCM. Of interest, a similar positive association of *HSPD1* (HSP60) expression with M2 markers expression was observed in human muscle tissues derived from the MoTrPAC meta-analysis following acute resistance exercise (**Figure 4C**), supporting the human relevance of our *in vitro* results.

**Figure 4.**
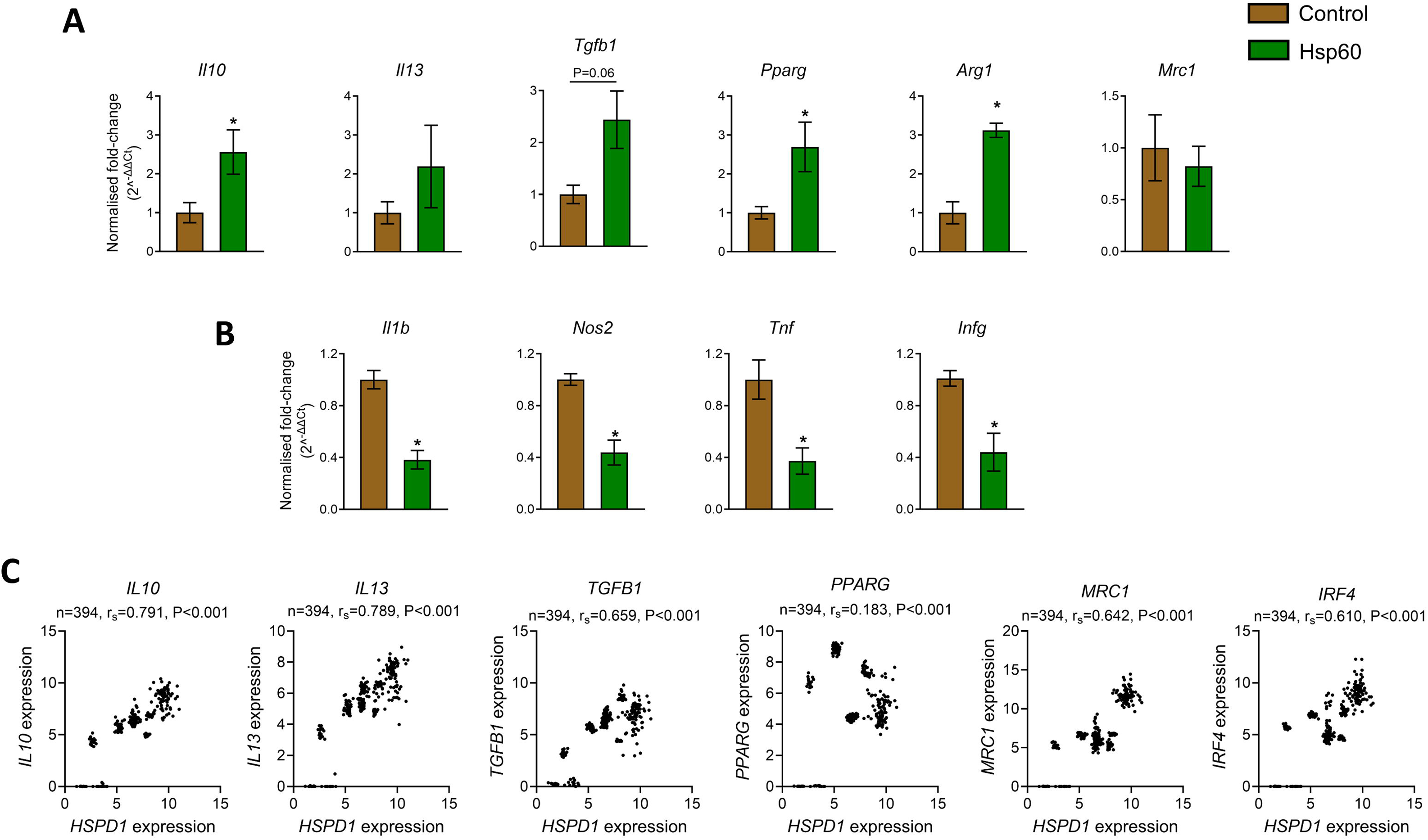
Treatment of BMDMs with HSP60 induces an anti-inflammatory phenotype. BMDMs were treated with 5mg/ml Hsp60 or vehicle (PBS; control group) for 24h (n = separate cell isolations from 5 mice). qRT-PCR analysis of BMDMs for (A) anti-inflammatory or (B) pro-inflammatory gene expression. Relative gene expression for each sample is expressed as fold-change *vs* the average value of the control group. Bars represent mean ± standard error of the mean (SEM) of each depicted group. Two-tailed Mann Whitney U-tests were used to calculate significance. **C)** Scatter-plots depicting the correlation between *HSPD1* (HSP60) and anti-inflammatory genes (M2 markers) in the human MotrPac meta-analysis dataset (available at: https://motrpac-data.org)* p˂0.05.

Collectively, these findings show that skeletal muscle Hsp60 is enriched following exercise *in vitro* and *in vivo*, and induces a transcriptional and bioenergetic switch in macrophages towards an anti-inflammatory phenotype in a paracrine/endocrine manner.

## Discussion

The beneficial effects of regular low-to-moderate intensity exercise in the prevention and alleviation of multiple non-communicable diseases is increasingly recognized (9). While some effects such as cardiovascular risk reduction have been long known, the molecular mechanisms underlying the anti-inflammatory effects of exercise remain largely unknown. Recently, macrophages, as key regulators of tissue microenvironment in multiple non-communicable disorders [reviewed in (25)], have gained significant attention as part of the exercise-induced anti-inflammatory responses. Exercise training in high-fat diet-fed obese rats has been shown to promote an M1-to-M2 switch of adipose tissue macrophages with concomitant reduction of proinflammatory gene expression (13). The pro-resolving properties of exercise were also shown in a model of wound-healing, where exercise promoted an M2 phenotypic switch of macrophages and accelerated wound healing (26). In line with this, an 8-week long program of moderate exercise promoted an M2-like phenotype of BMDMs in mice, showing dampened responses to immune challenge with LPS (27). This M2 switch was accompanied by changes in chromatin accessibility and expression of genes associated with inflammation and metabolism, as well as reliance of exercised BMDMs on oxidative phosphorylation (27). Of note, exercise training can also affect tissue-resident macrophages, as shown by the anti-inflammatory phenotypic switch of Kupffer cells in the liver accompanied by metabolic re-programming, which had significant protective effects against ischemia-reperfusion injury (28). Herein, we add to these previous studies showing that the exercise-induced, anti-inflammatory M2-like switch of macrophages may be mediated in a paracrine/endocrine manner by myotube secretome and specifically via the release of Hsp60.

The 60-KDa chaperonin protein (Hsp60) belongs to a highly conserved family of heat shock proteins that are essential for the folding and transportation of proteins (29, 30). Hsp60 is mainly located in the mitochondria, while intracellular translocations may occur under certain conditions (31–36). Unlike other chaperones (e.g., Hsp70, Hsp90) that share the α2-macroglobulin receptor as binding site, Hsp60 binds to a distinct receptor on macrophages, while Toll-like receptor (TLR)-4 is indispensable for downstream signaling and cytokine production (37). The release of Hsp60 or relocation on the cell surface was initially considered as a danger signal of stressed or damaged cells promoting proinflammatory responses via interactions with TLRs in innate immune cells, including macrophages (23, 38). In contrast, a recent study showed that extracellular Hsp60 induces an M2-like phenotype in skin macrophages and promotes wound healing in diabetic mice (24), which is in line with our results. Moreover, similar to what we observed in our mice that followed a program of aerobic exercise, increased Hsp60 was detected in the skeletal muscle and blood of mice following a 6-week endurance training program (39). More importantly, data from obese diabetic patients who followed a 3-month training program also showed significant increase of Hsp60 in subcutaneous adipose tissue (40). Future studies are warranted to examine the value of Hsp60 as a biomarker or therapeutic target in chronic inflammatory non-communicable disorders.

## Methods

### Cell Culture

C2C12 mouse myoblasts were used for the *in vitro* experiments of cyclic mechanical stretching. Cells were obtained from the American Type Culture Collection (Manassas, VA, USA) and were grown in Dublecco’s modified Eagle’s medium (DMEM) supplemented with 10% fetal bovine serum (FBS) and 1% penicillin/streptomycin in an atmosphere of 5% CO_2_ at 37°C. The medium was renewed every 2 days, and cells were split once they reached 70–80% confluence. For C2C12 differentiation after cells reached ∼ 80% confluence the cell media was changed to DMEM supplemented with 2% horse serum and 1% penicillin/streptomycin (differentiation medium). After the switch of the media, the multinucleated myotubes were easily distinguished and the cells reached their full differentiation potential.

### In vitro cyclic mechanical stretch

The Flexcell FX-5000 strain unit (Flexcell International) consists of a vacuum unit linked to a valve controlled by a computer program. Myotubes were cultured onto six-well Flexcell I flexible membrane dishes coated with type I Collagen (Flex I Culture Plates Collagen I; Flexcell International, Hillsborough, NC, USA) in high-glucose DMEM (Gibco) and were subjected to cyclic stretch produced by this computer-controlled application of sinusoidal negative pressure. C2C12 myotubes were either untreated (resting control) or exposed to a mechanical loading protocol of 2% elongation, 0.25 Hz frequency for 12 hours in DMEM containing 2.5% BSA as indicated in the figure legends (exercised), as previously described (17, 18). The myotube supernatant was then collected and stored at −80°C until further use.

### Culture of mouse primary bone marrow derived macrophages (BMDMs)

Bone marrow was isolated from the femurs and tibia of 6-8 weeks old C57Bl/6J male mice. Bone marrow cells were subsequently cultured for 7 days in DMEM full medium (10% FBS, 1% P/S) supplemented with 10 ng/ml M-CSF to induce macrophage differentiation. To examine the effect of the secretome of stretched myotubes on macrophages, isolated mouse BMDMs were cultured in the presence of C2C12 supernatant from stretched or resting (control) cells (1:1 dilution in DMEM containing 2.5% BSA and 1% Penicillin/Streptomycin) for 24 hours. In parallel experiments, the aforementioned culture conditions were repeated after pre-treatment of the cells with an OXHPOS inhibitor (30 nM IACS-010759, Selleckchem No. S8731) for 1h. In other experiments, BMDMs were treated with 5 μg/ml HSP60 (ab92364) in DMEM containing 2.5% BSA and 1% Penicillin/Streptomycin on the 7^th^ day of the culture for 24 hours.

### Mouse experiment

Eight-week-old C57Bl/6J male mice were used in this study. All animals were kept at a pathogen-free (SPF) controlled environment (22–26 °C temperature, 40–60% humidity, and 12 h light/dark cycle), with free access to water and a regular chow diet. All animal experiments were approved by the Region of Attica, Greece (536768, approved 16 September 2019).

Mice were divided into two groups: one group was subjected to an exercise protocol (exercise group) and the rest mice were not subjected to any exercise (sedentary group; control). Mice in the exercise group were subjected to treadmill running exercise 3 times/week for 9 weeks, with the first week being introductory at a lower speed and duration of exercise. On the first day of the introductory week, the mice were subjected to 10 min of walking/running at a speed increasing from 10 to 18 cm/s. The second day of the introduction consisted of 20 min of running with speeds up to 25 cm/s. During the third day of the introduction, the duration was increased to 30 min, with a speed of 25-35cm/s. The exercise protocol consisted of 30 min. treadmill running at a speed of 30-50 cm/s with no inclination of the treadmill (Panlab). After completing the exercise program, mice were euthanized and blood and quadriceps muscles were collected.

### RNA Isolation and qPCR

Total RNA was isolated from BMDMs using TRItidy G (AppliChem GmbH, Darmstadt, Germany) according to manufacturer’s instructions. The RNA concentration and purity were determined using a Nanodrop Spectrophotometer. RNA was reverse transcribed to complementary DNA (cDNA) using the PrimeScript™ RT Reagent Kit (Takara, Shiga, Japan). Gene expression was determined using the SsoFast EvaGreen Supermix (BioRad, Hercules, CA, USA) and specific primers for each target on an iQ5 Bio-Rad cycler system. The calculation of the relative gene expression was performed according to the ΔΔCt method and the eukaryotic translation elongation factor 2 (*Eef2*) served as housekeeping gene. For primer sequences please see ***Supplementary Table 1***.

### Quantification of circulating Hsp60 levels

To determine serum levels of Hsp60 a commercially available enzyme-linked immunosorbent assay (ELISA) kit was used (ab208344, Abcam, Cambridge, United Kingdom).

### Metabolomics

Upon cultivation, metabolite extraction from BMDMs took place in a 40:40:20 acetonitrile:methanol:water cell extraction solvent, as previously described (41). Raw mass spectrometry high-resolution data were acquired (70-1000amu) on a Triple TOF 6600 mass spectrometer operated on negative ion mode (ESI-) controlled through Analyst 1.8 software (AB Sciex, Warrington, UK). Electrospray ion source instrument parameters were optimized to Gas 1 60 (arbitrary unit), gas 250 (arbitrary unit), curtain gas 35 (arbitrary unit), source temperature 500℃, ion spray voltage −4500 V. A reversed phase ion pair chromatographic separation was employed to reduce metabolic complexity and improve mass detection (https://pubmed.ncbi.nlm.nih.gov/24861786/). Raw spectrometric data were processed with MultiQuant 3.0.3 software in parallel with an inhouse retention time/accurate mass library to extract metabolite peak areas. Raw spectrometric data were log2 scaled and normalised to median metabolite value on Excel spreadsheet. Log_2_-fold metabolite changes, ftest and ttest (two tails) were calculated based on binary group comparisons of interest. For each metabolite the analytical reproducibility was assessed based on coefficient of variation on the quality control samples. Significant metabolites were highlighted those which met all the following criteria |log(2) Fold Change|>0.58, QC CV<30% and p-value < 0.05. Metabolite validation results from Excel were imported to pathway analysis tool (inhouse built software) and pathway score values were assigned based on number of significant metabolites per metabolic pathway. The enrichment analysis in metabolomics data was applied using the up-regulated KEGG compound names (log2FC>0.58 and pvalues <0.05 and CV<30) and MetaboAnalyst 6.0 (https://www.metaboanalyst.ca/MetaboAnalyst/home.xhtml) platform, using the over-representation analysis in the Enrichment Analysis module.

### Seahorse Real-Time Cell Metabolic Analysis

7×10^4^ BMDM were plated in Seahorse XFe96 plates (V3-PS, TC treated, Agilent Technologies) and treated with exercise or control conditioned medium (50%) from C2C12 cells. Following activation, the cell medium was changed to XF RPMI (#103576-100, Agilent Technologies) supplemented with 2mM L-glutamine (#G7513), and 2mM sodium pyruvate (# P5280), with (mitostress) or without (glycostress) 10mM glucose (#G8270, Sigma-Aldrich), adjusted to pH 7.4 with the addition of 1M NaOH and incubated at 37 ℃, 0% CO2 for 1h. according to the manufacturer’s instructions. three sequential injections of 2μΜ oligomycin (#75351), 1μM FCCP (#C2920) and 2μΜ rotentone (#R8875)/ 2μΜ antimycin A (#A8674) were used for the mitostress protocol, while 10mΜ D-glucose (#G8270), 2μΜ oligomycin (#75351) and 50mΜ 2-DG (#D6134) were used for the glycostress protocol. The assay was performed in an XFe96 Seahorse analyzer (Agilent Technologies). Twelve cell-free wells served as the background for OCR and ECAR values. The basic respiratory and glycolytic parameters were calculated according to Agilent’s equations.

### Proteomic sample preparation

The secretome samples (supernatant of exercised or non-exercised myotubes), serum-free and conditioned, were subjected to the Sp3 method to bind in 50% ethanol, wash the bound proteins twice with 80% ethanol, and finally to digest them with a Trypsin/LysC protease mix (Promega) in 50 mM ammonium bicarbonate overnight on a thermomixer at 37°C. The next day, the peptide mixtures were recovered and dried down in a speed vac and their concentration was estimated by absorbance reading at 280 nm.

### LC-MS/MS Analysis

Samples were subjected to nLC-nESI MS/MS analysis in an Ultimate 3000 RSLC chromatographic system coupled to the HF-X Hybrid Quadrupole-Orbitrap mass spectrometer (Thermo Scientific). Peptides were loaded with 0.1% formic acid in water acetonitrile (Buffer A) to a Acclaim pepMap (100 μm internal diameter) trap column at 10 μL/min for 3.5 min. On chromatographic separation, peptides were loaded with 5 to 25% (65 min) to 40% (70 min) acetonitrile in 0.1% formic acid (Buffer B) on a 50 cm long (75 μm internal diameter) separation column at 300 nL/min. The buffer B was finally raised to 90% for 3 minutes and then the column was equilibrated for 15 minutes before the next sample injection.

Analysis was carried out in a data-dependent positive ion mode (DDA). The spray voltage was set to 1.9 kV with a capillary temperature of 270°C. Full scan MS were obtained on the Orbitrap analyzer at a 120,000 resolution with an Automatic Gain Control (AGC) set to 3×10^6^ and maximum injection time 50 ms. The 20 most abundant ions of each survey (from m/z 375 to 1500) were submitted to higher-energy collisional dissociation (HCD) of 27% NCE followed by new Orbitrap analysis at resolution of 15,000, using the following parameters: AGC at 8×10^3^ dynamic exclusion time of 60 s. The isolation window was set to 1.6 m/z. Peptide match was set to preferred and single charge ions were excluded. Data were obtained in technical duplicates using the Xcalibur software. The mass spectrometer was operated using the lockmass function on.

### Data analysis of the proteomic raw-files

The raw files were searched and the identified peptides and proteins were quantified using Label Free Quantitation (LFQ) in MaxQuant (version 1.6.14.0), using search against the mouse uniprot protein database (downloaded 09/03/2020). Search parameters included a molecular weight ranging from 350 to 5,000 Da, a precursor mass tolerance of 20 ppm, an MS/MS fragment tolerance of 0.5 Da, a maximum of two missed cleavages by trypsin, and methionine oxidation, deamidation of asparagine and glutamine and protein N-terminal acetylation were set as variable modifications. The protein and peptide false discovery rate (FDR) was set to 1%. The match-between-run and second peptides functions were enabled.

The statistical evaluation between the two secretome profiles was performed using the Perseus software (version 1.6.10.43). Proteins identified as “potential contaminants”, “reverse” and “only identified by site” were filtered out. The LFQ intensities were log2 transformed. Six biological replicates plus two corresponding technical replicates were grouped based on stretching or not. The proteins were filtered for a minimum of 5 valid values in at least one of the two groups. Missing values were imputed i.e. replaced by normal distribution, assuming that the corresponding protein is present in low amounts in the sample. A two-sided Student’s t-test of the grouped proteins was performed using permutation-based FDR correction <0.05 for truncation.

Proteins with statistically significant changes were kept and normalized through Z-scoring, before performing Euclidean hierarchical clustering of rows and columns (average). Venn diagrams were created with the R software environment. The mass spectrometry proteomics data have been deposited to the ProteomeXchange Consortium via the PRIDE partner repository with the dataset identifier PXD056231.

### Analysis of available transcriptomic datasets of exercised human muscle tissues

*HSPD1* expression levels were analyzed using the online interface of the MyoGlu dataset (https://exchmdpmg.medsch.ucla.edu/app/). Expression values were extracted from skeletal muscle samples in human data, categorizing the samples into overweight and normal-weight groups. The analysis included two distinct time points: A1 (Baseline, at rest) and A2 (Baseline, immediately after a 45-minute exercise session). The expression values were plotted to compare HSPD1 levels between these groups across the two conditions.

For the correlation analysis of HSPD1 with M2 markers, we used the data from the MotrPac meta-analysis dataset (https://motrpac-data.org). The dataset was downloaded, and the meta-analysis was conducted following the methodology described in the original publication. The analysis focused on the Acute exercise dataset, evaluating samples before and after resistance exercise. Correlation analysis was performed in R, employing the Spearman correlation method with exact = FALSE. Visualization of the correlation results was conducted using the ggplot2 package in R.

### Statistical analysis

Statistical analysis was performed using GraphPad Prism 9 or as indicated in individual methods above. Normality of distribution was tested by D’ Agostino Pearson and Shapiro-Wilk tests. Continuous variables are described as means ± standard error of the mean (SEM). Comparisons of continuous variables between 2 groups was performed by two-sided Mann Whitney U test unless otherwise specified in individual methods above. The level of statistical significance was set at p<0.05.

## Supporting information

Supplementary material for Exercise induces anti-inflammatory reprogramming in macrophages via Hsp60

## Study approval

All animal experiments were approved by the Region of Attica, Greece (536768, approved 16 September 2019).

## Data availability

The proteomic dataset has been submitted to ProteomeXchange via the PRIDE database (The data is currently private and can only be accessed with a single reviewer account that has been created. Data will be made publicly available after acceptance).

## Project Name

Exercise induces anti-inflammatory reprogramming in macrophages via Hsp60

## Project accession

PXD056231

