## Supplementary material for Exercise induces anti-inflammatory reprogramming in macrophages via Hsp60 for "Exercise induces anti-inflammatory reprogramming in macrophages via Hsp60"

Correspondence to:

| **Supplementary Table 1.** Sequences of the specific sets of primers used for RT-PCR analyses. | | |
|---|---|---|
| **Target Gene** | **5′-3′ Forward Primer Sequence** | **5′-3′ Reverse Primer Sequence** |
| *Il1b* | ATCCCAAGCAATACCCAAAG | GTGCTGATGTACCAGTTGGG |
| *Nos2* (iNOS) | ACCTTGTTCAGCTACGCCTT | CATTCCCAAATGTGCTTGTC |
| *Tnf* | AGCCCCCAGTCTGTATCCTTCT | AAGCCCATTTGAGTCCTTGATG |
| *Il10* | ATCGATTTCTCCCCTGTGAA | TGTCAAATTCATTCATGGCCT |
| *Pparg* | TGCTCAAGTATGGTGTCCATGAG | AGTGCATTGAACTTCACAGCAAA |
| *Il13* | ATTGGAGATGTTGGTCAGGG | GCATGGTATGGAGTGTGGAC |
| *Arg1* | GGAAAGCCAATGAAGAGCTG | AGCATCCACCCAAATGACAC |
| *Tgfb1* | AGCTGCGCTTGCAGAGATTA | AGCCCTGTATTCCGTCTCCT |
| *Mrc1* (CD206) | GTGCAGACAAAGGCTGCCGGA | CCGCCTTTCGTCCTGGCATGT |
| *Ifng* | CTGGAGGAACTGGCAAAAGG | CTGGACCTGTGGGTTGTTGA |
| *Eef2* | GATCAGATCCGTGCCATCATGGACA | GTAGAAGAGGGAGATGGCGGTGGA |

**Supplementary Figure 1**

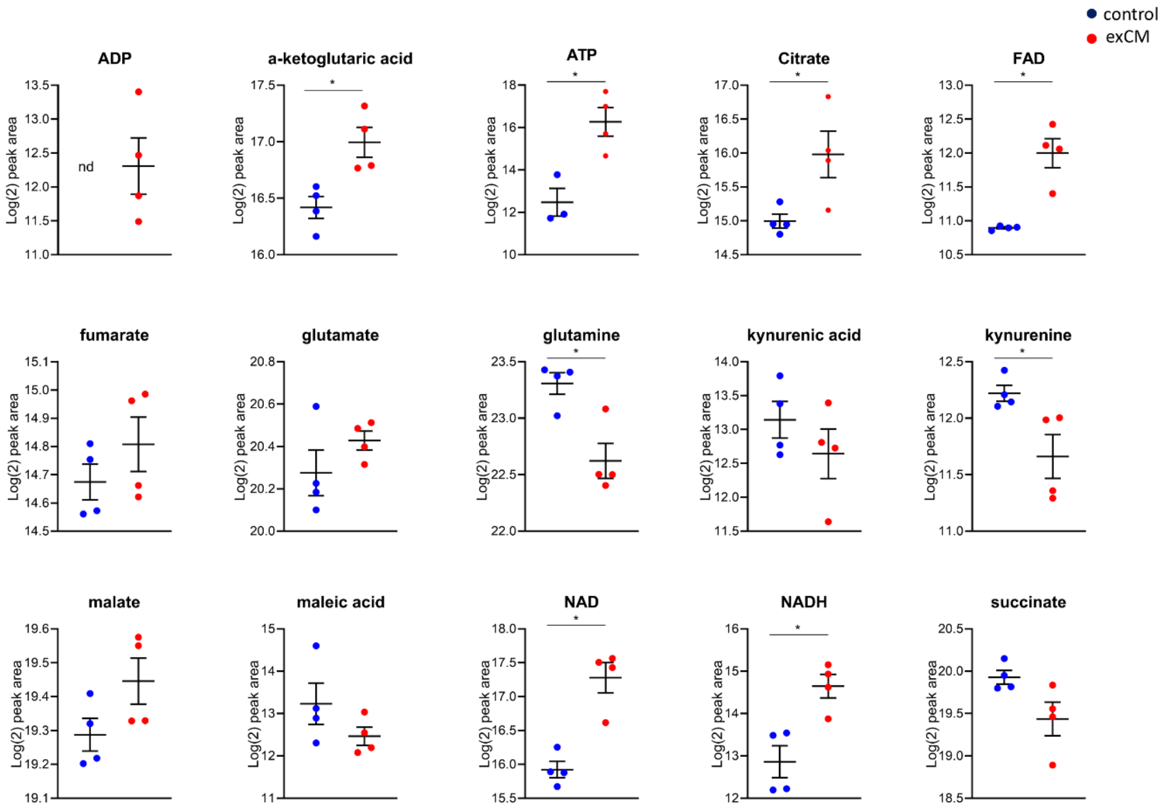


**Supplementary Figure 1. Levels of metabolites in macrophages following treatment with conditioned medium from exercised myotubes.** Fully differentiated C2C12 cells were subjected to 2% elongation for 12h using a cell stretcher. BMDMs treated with conditioned media (50%) from control (non-exercised) or stretched myotubes for 24h were subjected to metabolomic analysis (Non-exercised- vs exercised-treated BMDM from separate cell isolations from n=4 mice). Levels of selected metabolites, related to Figure 2A are shown here. Two-tailed independent samples t-test was used to calculate significance. nd: not detected, *P<0.05

**Supplementary Figure 2**

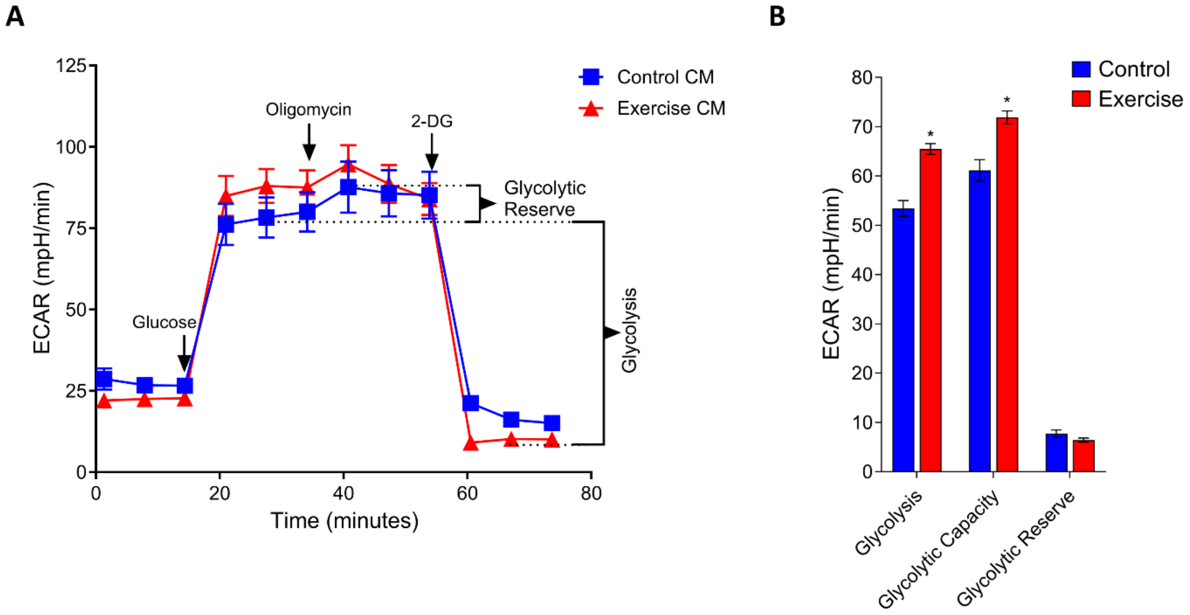


**Supplementary Figure 2.** Seahorse analysis of BMDMs for glycolysis (ECAR; extracellular acidification rate) bioenergetics. One representative experiment with 10 technical replicates per condition is shown in the line graphs. Bars represent mean ± SEM. Two-tailed Mann Whitney U-test was used to calculate significance. * P<0.05

**Supplementary Figure 3**

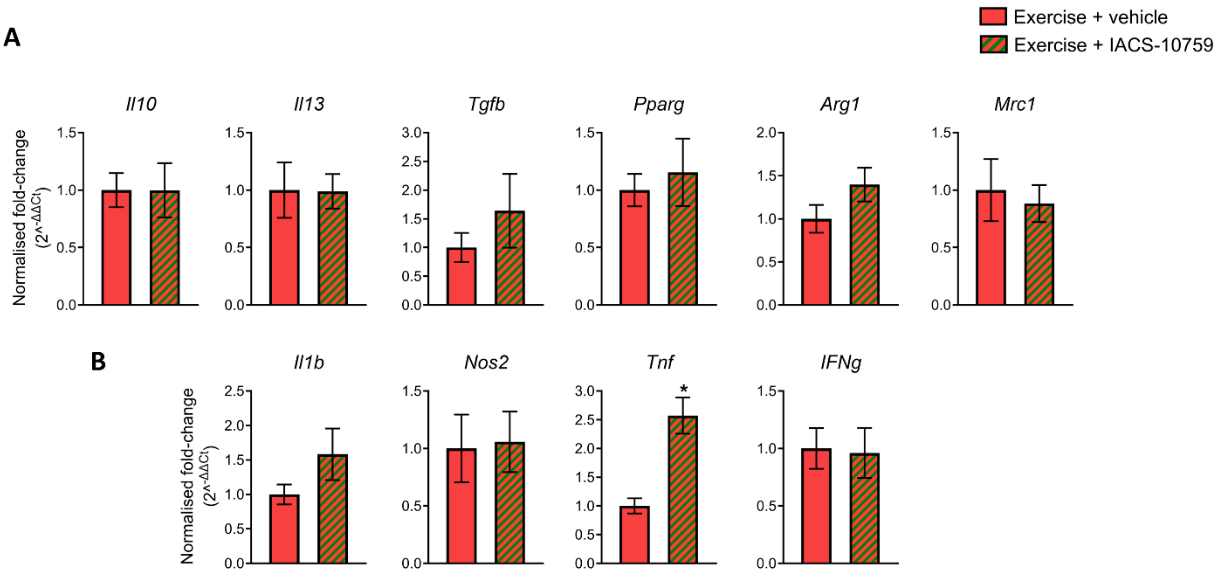


**Supplementary Figure 3. The effect of OXPHOS blockade on inflammatory gene expression in macrophages**. BMDMs were pre-treated with the OXPHOS inhibitor IACS-010759 for 1h (Exercise-anti-OXPHOS) or vehicle, and then treated with exercise-conditioned media (50%) from stretched myotubes for 24h (n = separate cell isolations from 5 mice). qRT-PCR analysis of BMDMs for (A) anti-inflammatory or (B) pro-inflammatory gene expression. Relative gene expression for each sample is expressed as fold-change *vs* the average value of the control group. Bars represent mean ± standard error of the mean (SEM). Two-tailed Mann Whitney U-test was used to calculate significance. * $P<0.05$
